# Event-related potentials reflect prediction errors and pop-out during comprehension of degraded speech

**DOI:** 10.1101/2020.03.24.005165

**Authors:** Leah Banellis, Rodika Sokoliuk, Conor J Wild, Howard Bowman, Damian Cruse

## Abstract

Comprehension of degraded speech requires higher-order expectations informed by prior-knowledge. Accurate top-down expectations of incoming degraded speech cause a subjective semantic “pop-out” or conscious breakthrough experience. Indeed, the same stimulus can be perceived as meaningless when no expectations are made in advance. We investigated the ERP correlates of these top-down expectations, their error signals, and the subjective pop-out experience in healthy participants. We manipulated expectations in a word-pair priming noise-vocoded speech task and investigated the role of top-down expectation with a between-groups attention manipulation. Consistent with the role of expectations in comprehension, repetition priming significantly enhanced perceptual intelligibility of the noise-vocoded degraded targets for attentive participants. An early ERP was larger for mismatched (i.e. unexpected) targets than matched targets, indicative of an initial error signal not reliant on top-down expectations. Subsequently, a P3a-like ERP was larger to matched targets than mismatched targets only for attending participants - i.e. a pop-out effect. Rather than relying on complex post hoc interactions between prediction error and precision to explain this apredictive pattern, we consider our data to be consistent with prediction error minimisation accounts for early stages of processing followed by Global Neuronal Workspace-like breakthrough and processing in service of task goals.

## Introduction

Prediction error minimisation accounts of perception propose that the brain seeks to minimise the mismatch between incoming sensory information and top-down expectations (Friston, 2010; Rao & Ballard, 1999). To successfully comprehend speech, a prediction error minimisation account argues that the listener must generate a set of expectations at multiple levels of representation to attempt to most accurately explain the auditory input (Paczynski & Kuperberg, 2012). Consistent with the role of expectations in speech comprehension, the amplitude of the N400 event-related potential (ERP) in response to the final word of a sentence increases with how unexpected that word is, given the context of the sentence (Kutas et al., 1984; Kutas & Federmeier, 2011). The N400 can, therefore, be characterised as an index of the amount of mismatch between a semantic prediction and the incoming stimulus - i.e. a semantic prediction error (Bornkessel-Schlesewsky & Schlesewsky, 2019; Paczynski & Kuperberg, 2012). Indeed, prediction error minimisation accounts of global brain function, such as free energy (Friston, 2010), propose that all evoked activity in the brain reflects this mismatch of prediction and stimulus, i.e. the prediction error (Clark, 2013; Rao & Ballard, 1999).

However, not all ERPs can be characterised parsimoniously within a narrow prediction error framework. For example, highly predictable events in rapid serial visual presentation (RSVP) that are associated with a subjective experience of conscious ‘breakthrough’ or ‘pop-out’ also elicit large ERPs from ∼300ms post-stimulus (i.e. the P300; (Donchin & Coles, 1988), while unpredictable events in the same stream of stimuli will elicit almost no evoked response (Bowman et al., 2013; Rohaut et al., 2015) - the opposite of what would be predicted if ERPs indexed prediction error only. To account for these apparently apredictive effects, prediction error minimisation accounts propose that attention increases the precision of predictions, and that prediction error is subsequently weighted by this precision (Kok et al., 2012). As a result, a range of ERP magnitudes, including late apredictive components such as the P300 in RSVP, can be explained as contributions from independently varying precision and prediction error (see also Heilbron & Chait, 2018).

The Global Neuronal Workspace is an alternative theory of neural processing that proposes that such apredictive evoked positivities with onsets ∼300ms post-stimulus reflect the ignition of a stimulus representation into a frontoparietal network for conscious access - whether that stimulus was or was not expected - while earlier ERPs index preconscious processes, including prediction errors (Dehaene et al., 1998; Sergent et al., 2005). Applying this model to speech comprehension, Rohaut et al. (2015) proposed a two-stage ERP profile, with an initial unconscious semantic prediction error in response to each word (the N400, typically onsetting around 200ms post-stimulus) and a late positive complex (LPC; in this case onsetting around 600ms post-stimulus) reflecting the ignition of meaning into conscious access. In support of this proposal, the N400 ERP has been observed in states of relative unawareness such as sleep, coma, and vegetative state (or unresponsive wakefulness syndrome) (Beukema et al., 2016; Ibáñez et al., 2006; Kotchoubey et al., 2005; Rämä et al., 2010) while the LPC has only been reported in conscious individuals, or in those who were conscious of and subsequently could report target words (Rohaut et al., 2015; Sergent et al., 2005; van Gaal et al., 2014).

We sought to test the proposal that early ERPs (<300ms post-stimulus) reflect preconscious prediction error processes and later ERPs (>300ms post-stimulus) reflect conscious access by investigating the comprehension of speech that has been degraded by noise-vocoding (Shannon et al., 1995). Consistent with the role of expectations in speech comprehension, a noise-vocoded speech stimulus that is entirely unintelligible to a naive listener can be rendered intelligible through priming - for example, by presenting a non-degraded version of the stimulus (i.e. a *matched* prime) immediately prior to the degraded stimulus (i.e. the target). When successfully primed in such a word-pair listening task, listeners experience a “pop-out” of the meaning of the degraded speech - i.e. subjective conscious access (Davis et al., 2005) - while an unrelated (or, *mismatched*) prime will not facilitate comprehension of the subsequent target.

Evidence suggests that successful comprehension of noise-vocoded speech requires attentional effort (Hervais-Adelman et al., 2012) and top-down expectations from frontal lobes (Sohoglu et al., 2012; Wild, Yusuf, et al., 2012). Therefore, we predict that distracted participants will be unable to use a prime word to generate a top-down expectation of the identity of an upcoming target, and will, therefore, neither exhibit a differential prediction error signal nor any subsequent apredictive evoked response to the target. Conversely, we expect that attentive participants will use the prime to generate top-down expectations of the identity of the degraded stimulus, and will therefore more readily comprehend the target. Consequently, and consistent with a two-stage Global Neuronal Workspace account (Rohaut et al., 2015), we expect attentive participants’ ERPs to exhibit an initial prediction error signal (i.e. larger evoked response to mismatched targets; cf. (Sohoglu et al., 2012) followed by an apredictive “pop-out” effect in which the ERP to the correctly-expected and comprehended targets is larger than that to the unexpected and predominantly unintelligible targets.

## Methods

### Participants

We recruited participants from the University of Birmingham via advertisement on posters or the online SONA Research Participation Scheme until we had achieved our desired sample size of 48 participants with usable data (24 per group; Median age = 20 years, Range = 18-33 years). Our inclusion criteria were: right-handed (from self-report), 18 to 35-years-old, monolingual speakers of British English, with no self-reported epilepsy, dyslexia, or uncorrected hearing impairment. We compensated participants with course credit or £10/hour of their time. The STEM Research Ethics Board of the University of Birmingham granted ethical approval for this study and written informed consent was completed by all participants. To achieve our final sample, we recruited 77 participants but rejected data from 29 participants due to an error of randomisation in the experimental code.

### Stimuli

A male first-language British English speaker recorded 288 monosyllabic English nouns taken from previous priming studies in our lab (see https://osf.io/m9ud5/ for the full stimuli list; mean length = 440ms, range = 264-657ms, sampling rate = 44100Hz). First, we randomly assigned the stimuli to one of four equal-sized lists (72 words per group), and manually swapped words across lists until the lists were matched on imageability, frequency (BNC), length in phonemes, and length in letters. Frequentist tests (ANOVAs) indicated no evidence that the four lists differed in word frequency (F(3, 284) = 0.233, *p* = 0.873), imageability (F(3, 231) = 0.779, *p* = 0.507), length in phonemes (F(3, 284) = 0.217, *p* = 0.885), or length in letters (F(3, 284) = <0.001, *p* = 1; see Supplementary Materials), and Bayesian equivalent tests (conducted with (JASP Team, n.d.); (Morey & Rouder, n.d.)) revealed strong evidence in favour of the null hypothesis for all variables (all BF10 = < .05; see Supplementary Materials). From these four matched lists, we created counterbalanced conditions across participants (see *Procedure*).

We manipulated the intelligibility of targets through noise-vocoding - originally a form of auditory distortion used to simulate the experience of hearing by the means of a cochlear implant (Shannon et al., 1995) (for scripts, see https://github.com/conorwild/matlab-audio-scripts/). Noise-vocoding retains the coarse temporal structure of the speech but reduces spectral clarity and fine temporal detail. The amplitude envelope from (approximately) logarithmically-spaced frequency bands is extracted and applied to bandpass-filtered noise of the same frequency band. Finally, the bands of envelope-modulated noise are recombined to create the final noise-vocoded stimulus (Davis et al., 2005). Using this method, we created six-band noise-vocoded versions of each stimulus to be used as the targets, and subsequently normalised each stimulus to its RMS ((Wild, Davis, et al., 2012; Wild, Yusuf, et al., 2012).

### Procedure

We randomly assigned each participant to be in the attentive or distracted group. In the attentive group, each trial began with the auditory presentation of the prime followed by the target with a stimulus onset asynchrony (SOA) of 1 second (see Figure 1). After 2.2-seconds, participants were cued by a tone (500Hz, 200ms duration) to rate the “noisiness” of the target on a scale of 1 (low) to 5 (high) via keyboard press. Following each rating, the next trial began after an inter-trial interval of between 1 and 2 seconds, selected randomly from a uniform distribution on every trial.

**Figure 1.**
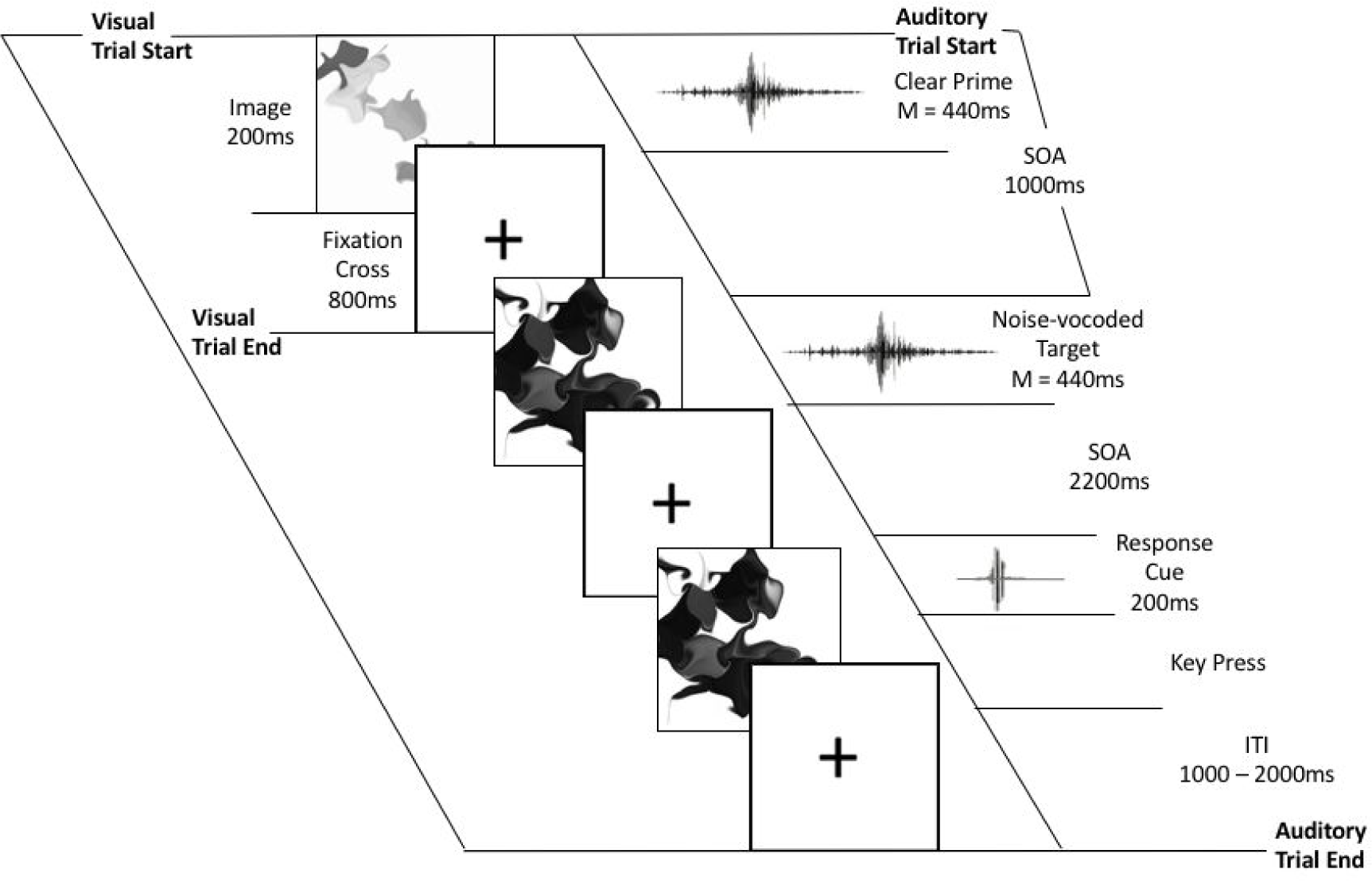
Schematic of event timing during the task.

In the distracted group, participants listened to the same auditory stimuli but did not complete the noisiness judgment task; instead, they performed a visual task (see below). Therefore, the timing of the auditory stimuli in the distracted group was identical to the attentive group, with the exception that the time between the onset of each target and the onset of the next trial was between 1 and 2 seconds, selected randomly from a uniform distribution on every trial.

While listening to the auditory stimuli, both attentive and distracted groups of participants watched a sequence of rapidly changing visual stimuli. However, only those in the distracted group were instructed to complete a task on the basis of the visual stimuli, and to ignore the auditory stimuli. The distraction task was a 1-back visual monitoring task, in which the sequence of visual stimuli was comprised of a series of images of ambiguous black shapes presented on a white background. Each ambiguous image was shown for 200ms with an 800ms fixation period between each image. For each participant, the order of images was randomized and the task was to press a key every time a repetition occurred (i.e. a 1-back task; 20% of trials). We subsequently calculated task accuracy to ensure that participants in the distracted group were distracted from the auditory stimuli by attending to the visual 1-back task. Participants in the attentive group watched the same visual stimuli, but were instructed to ignore them and to attend to the auditory stimuli only.

Upon completion of the above task, all participants completed a surprise recognition memory test, which included all 144 words from the mismatched condition of the word-pair priming task (i.e. 72 unrelated primes and 72 unrelated targets), as well as 72 new memory test items. We did not include the matched targets in the memory test as they were presented twice (as a clear prime and as a degraded target) and therefore cannot be compared to the unrelated targets or primes which were only presented once. In the memory test, each word was presented visually for 300ms, with a fixation point present for 2 to 3 seconds (selected randomly from a uniform distribution on each trial) between each word. We randomised the order of words for each participant. Participants made an old/new discrimination for each word on a 6-point remember-know scale, made up of the following responses; ‘definitely new, probably new, not sure, probably old, definitely old and remember’ (Ritchey et al., 2015). We reversed the scale for half of the participants to control for potential effects of response hand.

During the experiment, each participant heard all four word lists (see *Stimuli*), with each list comprising either the matched words, mismatched primes, mismatched targets, or new memory test items. For each of the 12 possible combinations of word lists for mismatched primes and mismatched targets, we manually ensured that there was no phonological, semantic, or associative overlap between the target and the prime. In total, there were 24 possible sets of stimuli to achieve full counterbalancing of lists. Therefore, across all participants, each word was heard an equal number of times in every possible condition.

### EEG pre-processing

We recorded EEG with a 128-channel Biosemi ActiveTwo system at a sample rate of 256Hz, with two additional electrodes recording from the mastoid processes. Offline, we digitally filtered the EEG signal between 0.5 and 40 Hz, segmented the data into epochs from 500-ms before the onset of the prime until 1000-ms after the onset of the target, re-referenced the data to the average of the mastoids, and baseline-corrected to the 200-ms pre-prime period. Unless otherwise stated, all offline pre-processing was performed with a combination of the Matlab toolbox EEGLAB (version 14.0.0b, (Delorme & Makeig, 2004)) and custom scripts. Note that all scripts are available online at https://osf.io/m9ud5/.

Artefact rejection proceeded in the following steps. First, we used an automated procedure, based on FASTER (Nolan et al., 2010), to identify and remove bad channels. Specifically, bad channels were those with absolute z-scores of >2.5 on any of the following measures: variance of voltage, mean correlations with other channels, and Hurst exponent. Across participants, a median of 7 channels were discarded (range 2-14). Second, we used an automated procedure, also based on FASTER (Nolan et al., 2010), to identify and remove trials with non-stationary artefacts. Specifically, a trial was bad if its absolute z-score was >2.5 on any of the following measures: mean range of voltages across channels, mean variance of voltages across channels, and the deviation of the trial average voltage from the average voltage across all channels. Third, we conducted Independent Component Analysis (ICA) of the remaining data (EEGLAB’s *runica* algorithm) and used the toolbox ADJUST (Mognon et al., 2011) to automatically identify and remove components with the expected spatial and temporal features of blinks, eye-movements, and generic discontinuities. Next, we interpolated any previously removed channels back into the data. Finally, trials with artefacts that had not been effectively cleaned by the above procedure were identified with visual inspection and discarded. After these pre-processing steps, a median of 65.5 trials contributed to the match condition (range: 39-71) and a median of 66 trials contributed to the mismatch condition (range: 37-71). Prior to analysis, all data were re-referenced to the average of all channels.

For our subsequent memory ERP contrasts, we only included data from the attentive group as recognition memory was not significantly greater than chance for the distracted group. We also excluded those participants who contributed fewer than 12 trials to either of the two categories (i.e. targets that were subsequently remembered [hits] versus targets that were subsequently forgotten [misses]) and those whose recognition memory was not greater than zero, resulting in a subgroup of 13 participants (hits: median 21, range 13-36; misses: median 26, range 12-49).

### EEG / MRI co-registration

The electrode locations of each participant were recorded relative to the surface of the head with a Polhemus Fastrak device using the Brainstorm Digitize application (Brainstorm v. 3.4; Tadel et al., 2011) running in Matlab (Mathworks). Furthermore, on a separate day, we acquired a T1-weighted anatomical scan of the head (nose included) of each participant with a 1mm resolution using a 3T Philips Achieva MRI scanner (32 channel head coil). This T1-weighted anatomical scan was then co-registered with the digitised electrode locations using Fieldtrip (Oostenveld et al., 2011).

### Sensor analyses: ERPs

Prior to analysis, we calculated participant-wise average ERPs for each condition separately using the robust averaging method of SPM12 (default params) that iteratively down-weights outlier voltages across trials. As recommended in the SPM12 documentation, the subsequent average ERPs were then low-pass filtered at 20Hz, and baseline-corrected to the 200ms prior to the onset of the target.

Analyses of ERPs proceeded in two stages, and in a similar way to (Sohoglu et al., 2012). First, we calculated the global field power (Skrandies, 1990) of the grand average of all trials (i.e. both conditions together) to identify time-windows of interest. Global field power (GFP) is the root mean square of average-referenced voltages, and is a principled means of identifying component peak latencies from an orthogonal contrast (Skrandies, 1990). We then identified a time-window around each peak by inspecting the global dissimilarity (Skrandies, 1990) – the mean of the root mean square of voltage differences between consecutive time-points, after the data have been scaled by the global field power. Deflections in the time-course of global dissimilarity therefore suggest boundaries between scalp topographies. On this basis, we selected the following ERP topographies: 137-ms – 207-ms, 211-ms – 246-ms, 250-ms – 371-ms, 375-ms – 547-ms, 551-ms – 648-ms, and 652ms – 707-ms (Figure 2).

**Figure 2.**
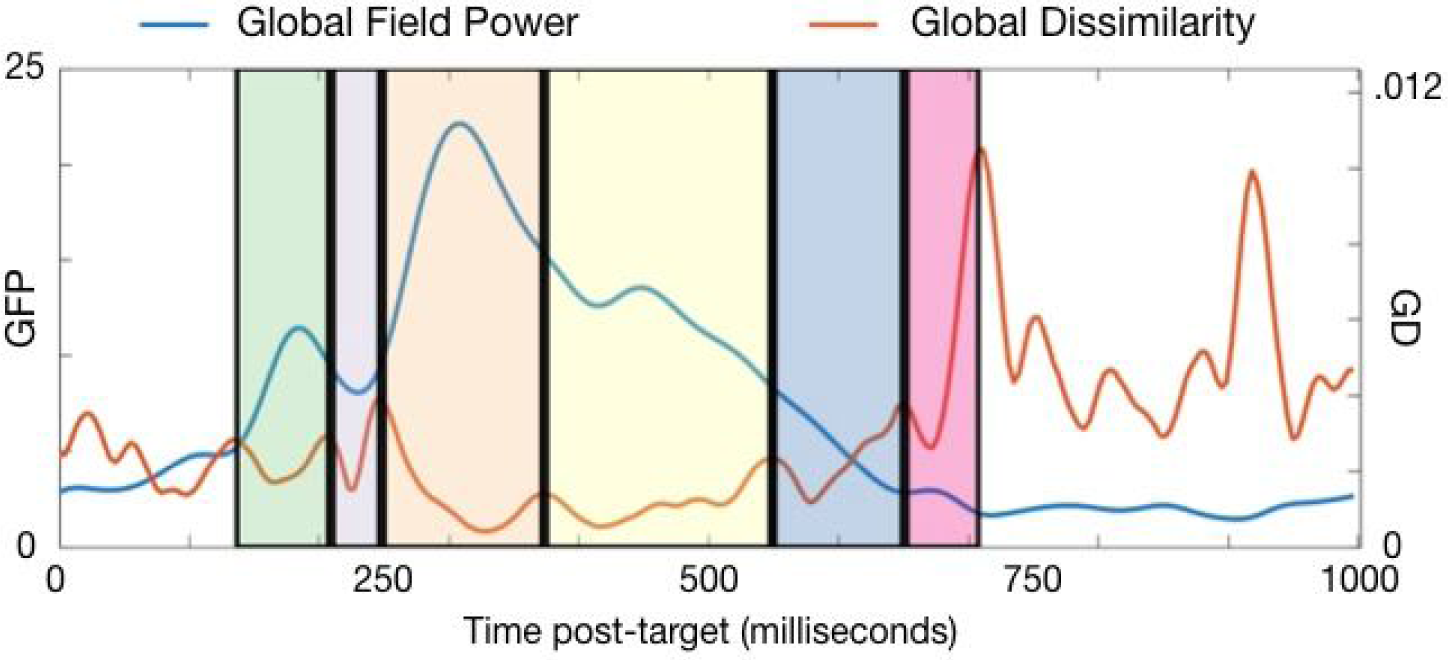
Global Field Power and Global Dissimilarity in the post-target window, with the time-windows of interest highlighted.

To minimise the number of comparisons in post-target time-windows, we only investigated the main effects of target type (i.e. matched versus mismatched, averaged across attention group) and attention (i.e. attention versus distraction, averaged across target type) when an interaction contrast produced no significant clusters (i.e. the difference between matched targets and mismatched targets between attention groups). Where a significant interaction cluster was observed, we tested for simple effects with paired samples t-tests of data averaged within the electrodes that contributed to the interaction cluster. Furthermore, within each significant interaction and main effect cluster, we investigated subsequent memory effects (hits versus misses) with paired samples t-tests of data averaged within the electrodes that contribute to each cluster.

ERPs (or difference ERPs between two within-subject conditions) within each time-window of interest were compared with the cluster mass method of the open-source Matlab toolbox FieldTrip (version 20160619, Oostenveld et al., 2011). First, for each participant x condition we averaged the voltages at each electrode within the time-window of interest. Next, a two-tailed t-test (dependent samples for interaction and main effect of target type; independent samples for main effect of attention) between conditions was conducted at each electrode. Spatially adjacent t-values with p-values passing the threshold (alpha = .05) were then clustered based on their spatial proximity. Clusters were required to involve at least 4 neighbouring electrodes, with an electrode’s neighbourhood defined as all electrodes within approximately 4-cm on a template head (median number of neighbours: 11; range: 2-16). A second non-parametric step corrects for multiple comparisons by conducting 1000 Monte Carlo randomisations of the above method (shuffling condition labels) to estimate the probability of the observed cluster under the null hypothesis (Maris & Oostenveld, 2007). We applied a cluster alpha threshold of .025 as we were testing for both positive and negative effects.

### Sensor analyses: Bayesian tests

When the above sensor analyses failed to find support for an interaction between target type and attention (i.e. the difference of differences) but did find evidence of a main effect, we used Bayesian equivalent t-tests to test the sensor data for evidence in support of the null hypothesis. Specifically, at each electrode we calculated a Jeffrey-Zellner-Siow Bayes factor (JZS-BF), implemented with an open-access script (https://github.com/anne-urai/Tools/tree/master/stats/BayesFactors). A JZS-BF > 3 is considered to be substantial evidence in support of the hypothesis being tested. While this approach does not take into account spatial clustering, as in the sensor analyses above, it does allow us to qualitatively inspect the spatial distribution of evidence in support of the null hypothesis across the head.

### Source estimation

We performed source estimation using EEG data and individual electrode locations from 48 participants. The analyses were completed using subject-specific T1-weighted anatomical MRI scans for 39 participants and template T1-weighted MRI images (provided by the Matlab toolbox FieldTrip) for the remaining nine participants due to issues with T1 data collection and image quality.

From the subject-specific T1-weighted anatomical scans, individual boundary element head models (BEM; four layers) were constructed using the ‘dipoli’ method of the Matlab toolbox FieldTrip (Oostenveld et al., 2011). Individual electrode locations were aligned to the surface of the scalp layer extracted from the segmented T1-weighted anatomical scans using fiducial points and head shape as reference points. The alignment of electrodes and scalp surface were further visually inspected to detect potential deviations and, where necessary, small manual corrections were applied.

As we required single-trial data to estimate the sources of the ERP effects, we used the pre-processed sensor-level data prior to the robust averaging step described above. We defined trials as time windows from −500ms to 1900ms relative to the prime, and baseline corrected the EEG data using the time window [−200ms – 0ms] relative to target presentation. Before the direct statistical comparison, we balanced the number of trials between conditions by randomly removing trials from the condition (discarded trials: median = 2, range = 0-13) with more data until both datasets had the same number of trials (median = 130, range = 74-136).

Our source estimation followed the analysis approach described in Popov et al. (2018). Therefore, the data was first filtered between 1 and 40 Hz, using a firws filter as implemented in the *ft_preprocessing* function of Fieldtrip (using default parameters). Additionally, to mitigate the confounding influence of correlated activity in the auditory cortices (i.e. from binaural stimulation) on LCMV beamformer source analysis, we calculated the surface Laplacian of the data and leadfields as in Murzin et al. (2013). Thus, the scalp current density of the data was calculated and the covariance matrix was estimated using a time window from −500ms to 1900ms. A common spatial filter (including trials of both conditions) was computed using an LCMV beamformer (inputting the surface Laplacian transformed leadfield) (Robinson, 1999; Van Drongelen et al., 1996; Van Veen et al., 1997). Specific beamformer parameters were chosen based on the approach used by Popov et al. (2018) including a fixed dipole orientation, a weighted normalisation (to reduce the centre of head bias), as well as a regularisation parameter of 5% to increase the signal to noise ratio. This common spatial filter was then used for source estimation. The dipole moments of both conditions were extracted in the post-stimulus time windows that showed significant clusters at the sensor level (time window 1: 137ms – 207ms; time window 2: 211ms – 246ms; time window 3: 250ms – 371ms; time window 4: 551ms – 648ms), and their absolute values were averaged over time points to obtain one average value per grid point (virtual electrode) and time window of interest. For clear visualisation of the foci of our source estimates, we calculated t-tests at each virtual electrode and thresholded the subsequent t-images at p<.05 (see Supplementary Materials and Sokoliuk et al. (2019), for further validation of the method).

## Results

### Speech intelligibility

Attentive participants rated the mismatched targets as noisier (Median = 4, Range 1-5) than the matched targets (Median = 3, Range 1-4), despite the stimuli being physically distorted to the same level. This difference was significant in a Wilcoxon Signed-Rank Test (W = 232, *p* = <.001).

### Recognition Memory: Discrimination (d’)

Discrimination (d’) was calculated as the z-transformed proportion of hits minus the z-transformed proportion of false alarms (Haatveit et al., 2010). The proportion of hits and false alarms were transformed using the inverse of the standard normal cumulative distribution. All ‘probably old’, ‘definitely old’, and ‘remember’ responses to old items were considered a hit, while the same responses to new items were considered false alarms.

A two-way mixed ANOVA with factors of word type (clear prime; degraded target; both only heard once by each participant) and attention (attentive; distracted) revealed significant main effects of word type (F(1,46) = 30.243, p = ≤ .001, partial n2 = .397) and attention (F(1,46) = 8.714, p = .005, partial n2 = .159), and a non-significant interaction (F(1,46) = 0.528, p = .471, partial n2 = .011). A Bayesian equivalent mixed ANOVA revealed considerable evidence for a model containing main effects of both word type and attention (BF=69083 relative to a null model), which itself was 2.687 times more likely given the data than a model containing both main effects and an interaction term. These results reflect the participants’ more accurate memory for clear primes than degraded targets and the higher memory accuracy in the attentive group than the distracted group.

One-Sample T-Tests determined that d’ for both clear primes and degraded targets were significantly different from zero (i.e. above chance) for attentive participants (Clear: Mean d’ = 0.457, SD = 0.309, t(23) = 7.245, p = ≤ .001; Degraded: Mean d’ = 0.194, SD = 0.218, t(23) = 4.368, p = ≤ .001), while memory for either item type was not significantly different from zero for distracted participants (Clear: Mean = 0.132, SD = 0.515, t(23)= 1.251, p = .223; Degraded: Mean = −0.070, SD = 0.393, (t(23) = −0.869, p = .394), suggesting that the distraction task effectively suppressed processing of the auditory stimuli. Bayesian equivalent T-Tests indicated considerable evidence for better than chance memory for attentive participants (Clear: BF_10_ = 71004; Degraded: BF_10_ = 131), and anecdotal evidence for chance-level memory performance for distracted participants (Clear: BF_10_ = 0.430; Degraded: BF_10_ = 0.302).

### Recognition Memory: Recollection & Familiarity

To estimate the level of processing that the clear and degraded words received, we calculated separate measures of Recollection (i.e. explicit contextualised memory of the event) and Familiarity (i.e. memory without context) from the recognition memory judgments for each participant (Atkinson & Juola, 1974; Yonelinas et al., 1997). Specifically, Recollection scores were calculated by: (Rold – Rnew)/(1-Rnew), with Rold reflecting the proportion of old items given a Remember response by the participant, and Rnew reflecting the proportion of new items given a Remember response. Familiarity was calculated by: (Fold/(1-Rold)) - (Fnew/(1-Fnew)), with Fold reflecting the proportion of old items given a ‘definitely old’ or ‘probably old’ response by the participant, and Fnew reflecting the same responses to new items (Ritchey et al., 2015).

Two two-way mixed ANOVAs with factors of word type (clear prime; degraded target; both only heard once by each participant) and attention (attentive; distracted) revealed significant main effects of word type on both Recollection and Familiarity estimates (F(1,46) = 13.287, p ≤ .001, partial n2 = .224, and F(1, 46) = 14.533, p ≤ .001, partial n2 = .240, respectively), reflecting higher scores for clear primes than for degraded targets. No other main effect or interaction was significant (all ps>.127). Bayesian equivalent ANOVAs similarly concluded that there was substantial evidence for models containing a main effect of word type for both Recollection (BF_10_ = 43.703, BF_inclusion_ = 60.743) and Familiarity (BF_10_ = 64.090, BF_inclusion_ = 42.913).

One-sample t-tests identified significantly different from zero measures of Recollection for clear primes (t(23) = 6.127, p ≤ .001, BF_10_ = 6538.041) and degraded targets (t(23) = 2.499, p = .020, BF_10_ = 2.720) in the attentive group, while Familiarity was not different from zero (Clear: t(23) = −0.009, p = .993, BF_10_ = 0.215; Degraded: t(23) = −1.899, p = .070, BF_10_ = 0.998). In the distracted group, neither Recollection nor Familiarity were significantly different from zero for either word type (Recollection Clear: t(23)= 0.245, p = .809, BF_10_ = 0.221; Recollection Degraded: t(23) = −1.003, p = .326, BF_10_ = 0.337; Familiarity Clear: t(23)= −1.452, p = .160, BF_10_ = 0.542; Familiarity Degraded: t(23) = −2.042, p = .053, BF_10_ = 1.247).

### Event-related Potentials

#### Interaction effects

We observed an interaction between target type and attention in the 250-371-ms time-window post-target only (cluster p=.011) with estimated generators in right middle temporal gyrus and right fusiform gyrus (Figure 3). Follow-up simple effects tests indicated greater positivity within this cluster for matched targets relative to mismatched targets during auditory attention only (t(23)=2.755, p=.011, two-tailed, BF10=4.376; Figure 3D). While the mean voltages in this time-window exhibited the opposite pattern in the distracted group - i.e. greater positivity to mismatched targets - this difference did not pass our significance threshold (t(23)=-1.869, p=.074, two-tailed; Figure 3D) and a Bayesian equivalent analysis found only anecdotal evidence in favour of the null hypothesis in this contrast (BF10=.955). A subsequent memory contrast within this cluster indicated greater positivity for mismatched targets that were subsequently remembered (hits) relative to mismatched targets that were subsequently forgotten (misses) in the attentive group, although this effect was only weakly significant in a t-test (t(12)=2.185, p=.049, two-tailed; Figure 3E) and a Bayesian equivalent indicated that the evidence was only anecdotal (BF10=1.628). No clusters were formed in any other time-window for the interaction contrast. We therefore examined the main effects below.

**Figure 3.**
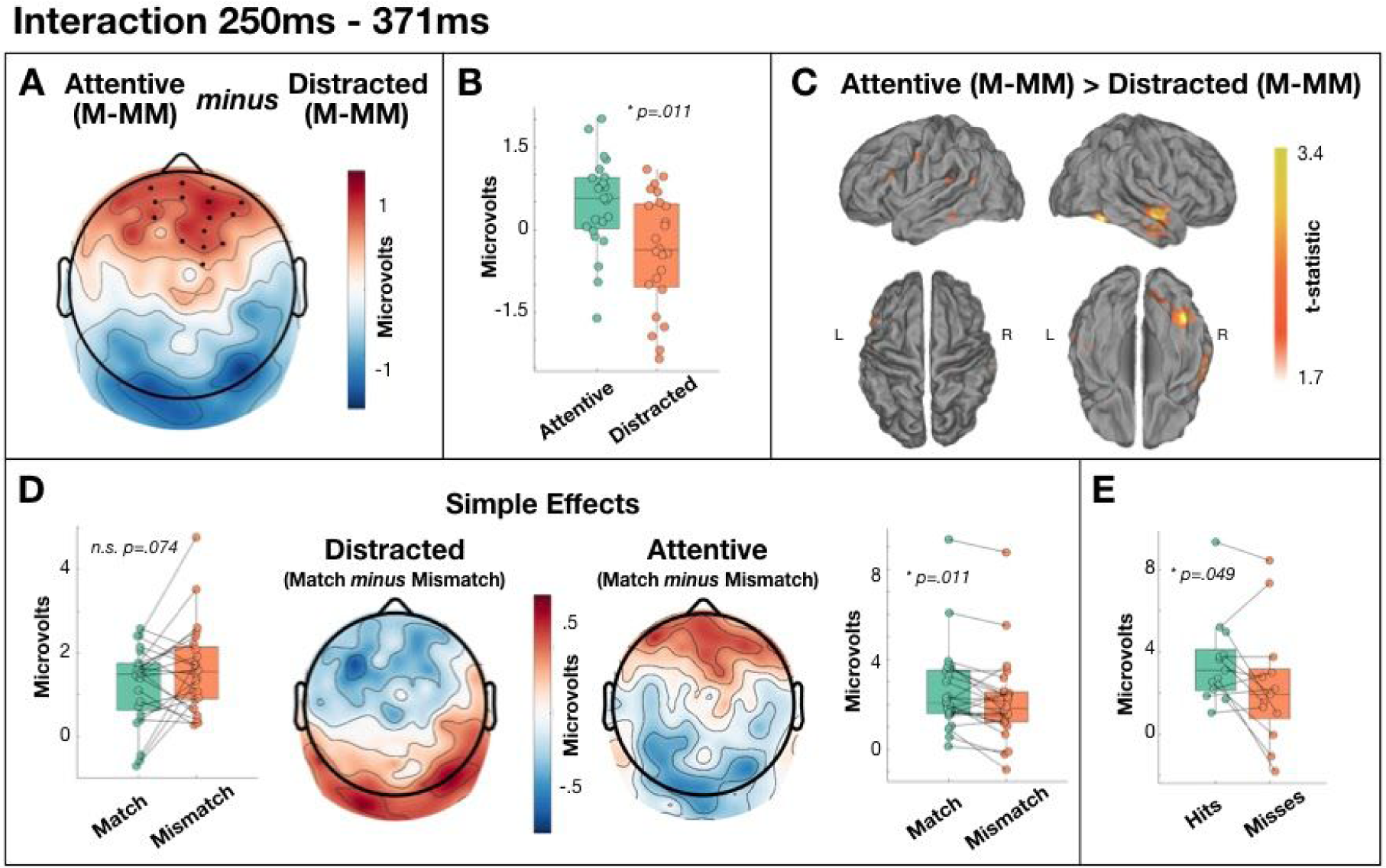
Interaction between target type and attention from 250ms-371ms. [A] Scalp distribution of the significant difference in the effect of target across attention conditions. Electrodes contributing to the cluster are marked. M-MM: match minus mismatch. [B] Single-subject mean difference voltages (difference between match and mismatch) within the significant cluster. [C] Estimated sources of the attentive effect of target in right middle temporal gyrus and right fusiform gyrus (relative to the distracted effect of target). [D] Analysis of the simple effects showing qualitatively different topography across attention groups, and a significant effect of target type in the attentive group only. [E] Subsequent memory effect within the interaction cluster.

#### Main effects

We observed a dipolar main effect of attention in the 137-207ms time-window, with greater frontal positivity (cluster p=.009) and greater posterior negativity (cluster p=.010) for attentive participants relative to distracted participants (Figure 4). Our source analyses estimated this effect to be generated primarily within right superior frontal lobe, overlapping with right premotor cortex. A subsequent memory contrast within each cluster indicated significantly larger ERPs for subsequent hits relative to subsequent misses in the attentive group, with the Bayes factor in the frontal cluster indicating substantial evidence for a subsequent memory effect (positive frontal cluster: t(12)=2.657, p=.021, BF10=3.195; negative posterior cluster: t(12)=-2.320, p=.039, BF10=1.963).

**Figure 4.**
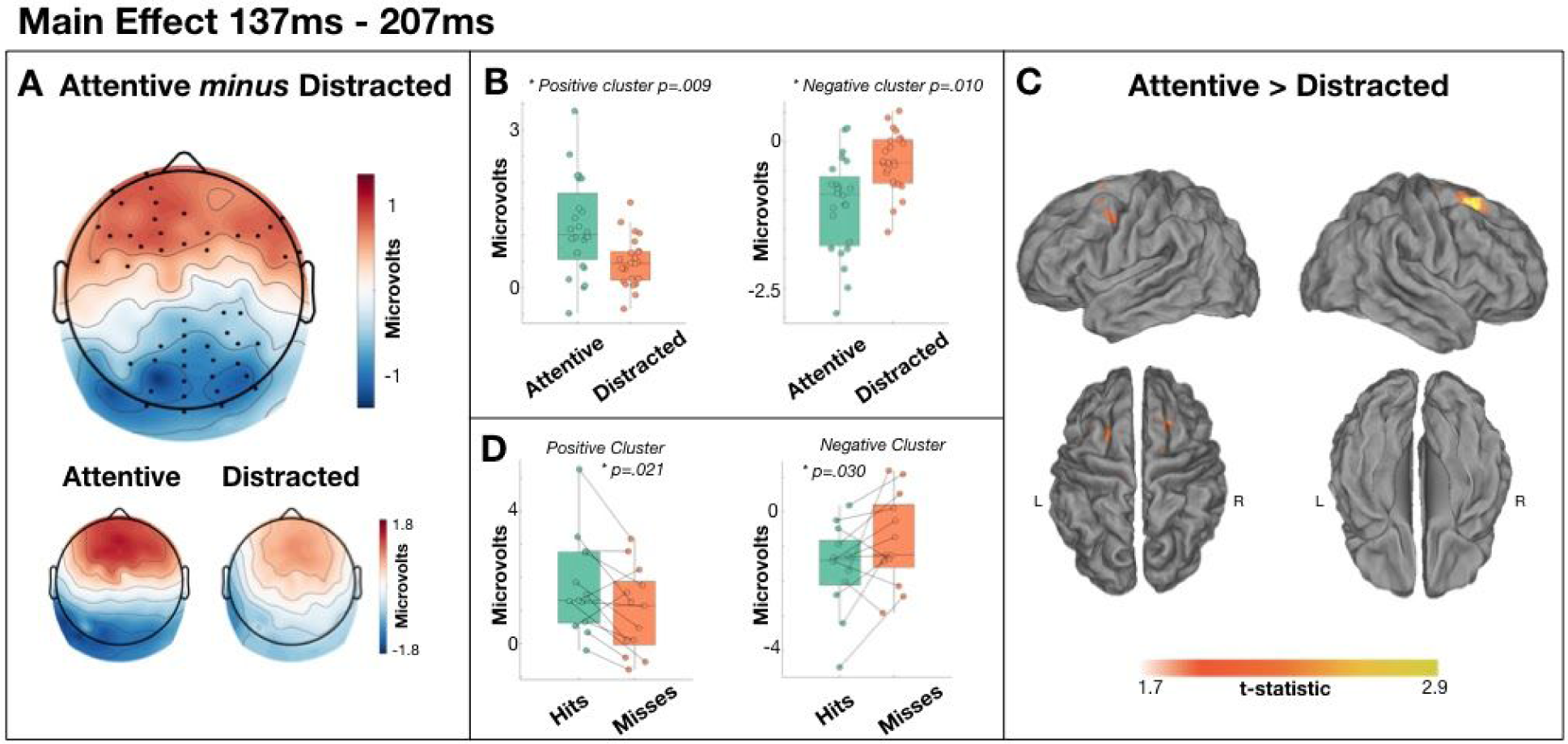
Main effect of attention from 137ms - 207ms. [A] Scalp distribution of the significant effect. Electrodes contributing to the clusters are marked. [B] Single-subject mean voltages within each significant cluster. [C] Estimated sources of the main effect within right superior frontal lobe. [D] Subsequent memory effects within each cluster in the attentive group.

We also observed a dipolar main effect of target type in the 211-246ms time-window with a larger left frontocentral positivity to mismatched targets than to matched targets (cluster p=.023) and a larger right temporal negativity to mismatched targets than to matched targets (cluster p=.013; Figure 5 A-C). Source analyses estimated this effect to be primarily generated within left supramarginal gyrus and right insula. Both Frequentist and Bayesian t-tests indicated no compelling evidence of subsequent memory effects in the attentive group in either cluster (positive cluster: t(12)=-1.764, p=.103, BF10=0.939; negative cluster: t(12)=1.964, p=.073, BF10=1.210).

**Figure 5.**
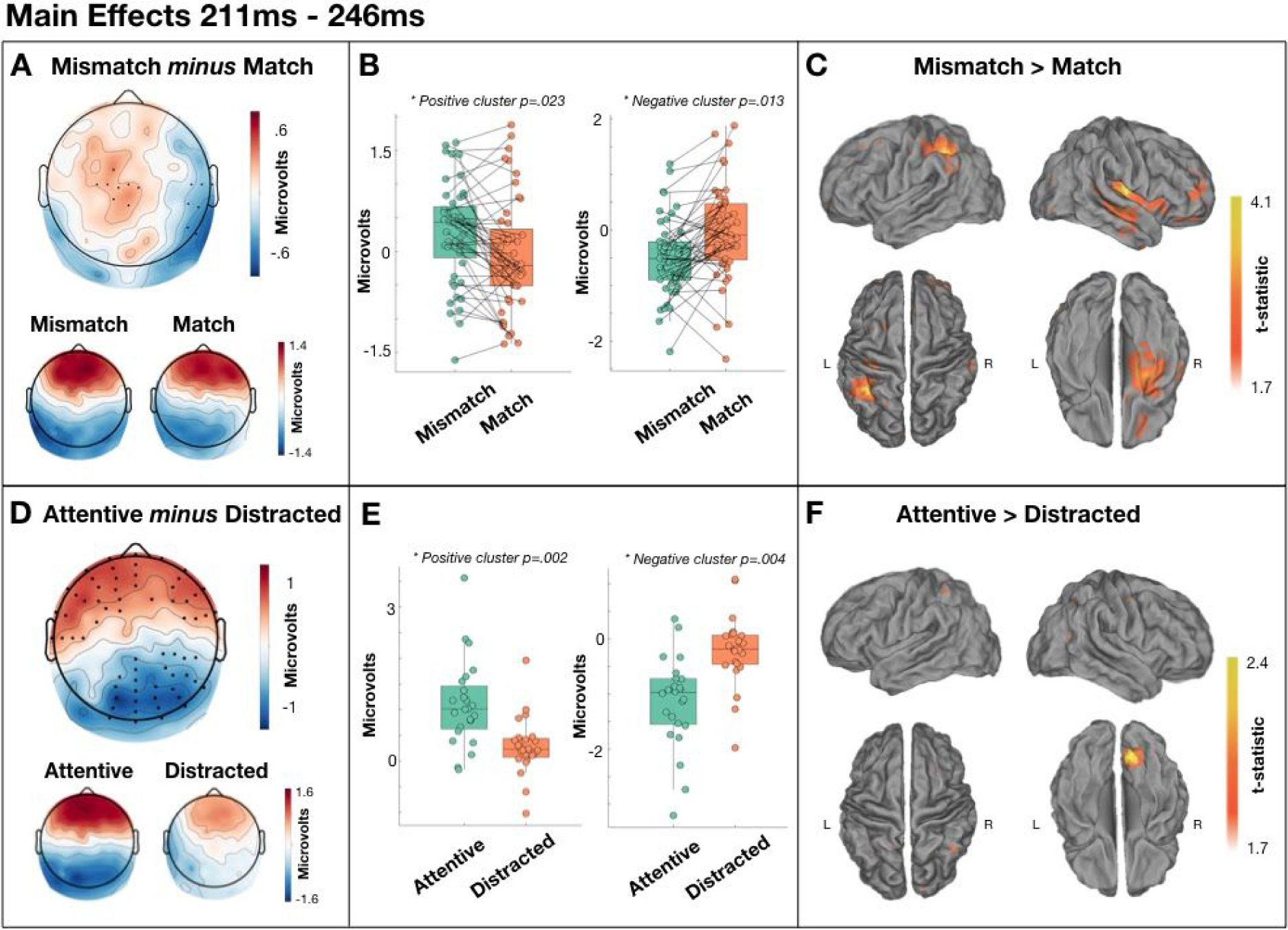
Main effects of target type and attention from 211ms - 246ms. [A] Scalp distribution of the significant effect of target type. Electrodes contributing to the clusters are marked. [B] Single-subject mean voltages within each significant cluster. [C] Estimated sources of the main effect within left supramarginal gyrus and right insula. [D] Scalp distribution of the significant effect of attention. Electrodes contributing to the clusters are marked. [E] Single-subject mean voltages within each significant cluster. [F] Estimated sources of the main effect within right visual cortex.

In the same time-window (211-246ms), we also observed a main effect of attention, with greater frontal positivity (cluster p=.002) and greater posterior negativity (cluster p=.004) in the attentive group relative to the distracted group, with estimated generator in right visual cortex (Figure 5 D-F). As with the effect of target in this time-window, both Frequentist and Bayesian t-tests agreed that there is no evidence of subsequent memory effects in either cluster (positive cluster: t(12)=1.557, p=.145, BF10=0.734; negative cluster: t(12)=-1.443, p=.175, BF10=0.647).

In the 551-648ms time-window, we observed an effect of target type, with a larger centroparietal negativity to mismatched targets than matched targets (cluster p=.019) estimated to be generated in the right posterior superior temporal gyrus (Figure 6). The subsequent memory contrast in this cluster failed to reach our significance threshold, and the Bayesian equivalent similarly concluded only anecdotal evidence in favour of the hypothesis (t(12)=2.158, p=0.052, BF10=1.568).

**Figure 6.**
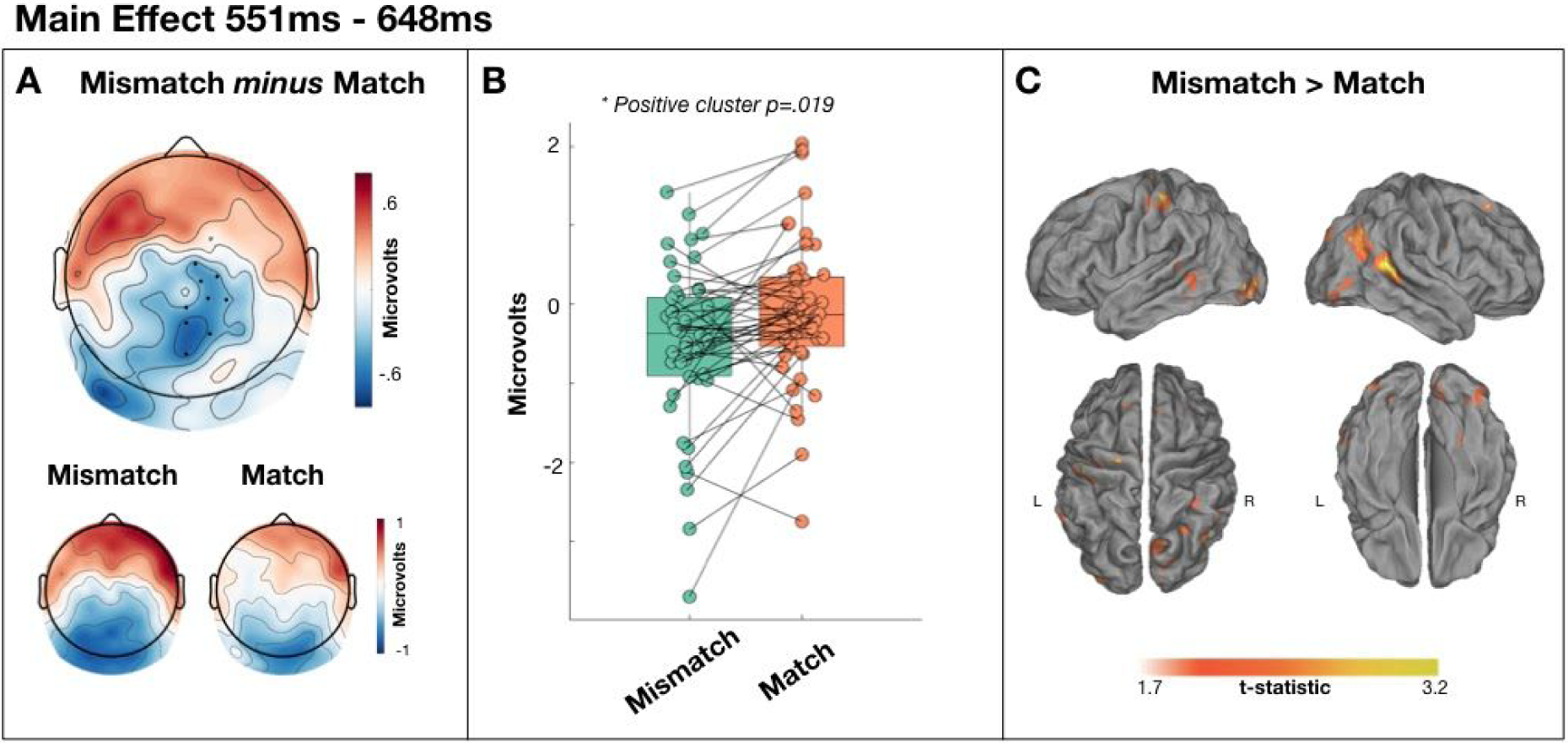
Main effect of target type from 551ms - 648ms. [A] Scalp distribution of the significant effect. Electrodes contributing to the cluster are marked. [B] Single-subject mean voltages within the significant cluster. [C] Estimated sources of the main effect within right posterior superior temporal gyrus.

The main effect of target in the 137-207ms and 652-707ms time-windows did not pass our significance threshold (p=.048 and .042 respectively; alpha=.025). The main effect of attention in the 375-547ms, and 652-707ms time-windows also did not pass our significance threshold (p=.047 and .082 respectively; alpha=.025). No clusters were formed in any other contrast or time-window.

## Discussion

Consistent with our hypothesis and previous research, repetition priming enhanced the perceptual intelligibility of the degraded targets (Davis et al., 2005; Hervais-Adelman et al., 2012; Sohoglu, 2014; Wild, Yusuf, et al., 2012). This result is, independently, evidence for the importance of prior knowledge (or expectation) for generating a conscious experience of comprehending degraded speech - i.e. a “pop-out”. Furthermore, consistent with a proposed two-stage ERP profile of auditory processing (Rohaut et al., 2015), we observe two dissociable ERP effects.

First, and contrary to some arguments of attentional enhancement of prediction errors (Auksztulewicz & Friston, 2015), we observe an early predictive signal (211-246ms) - i.e. larger for unpredicted words than for predicted words - that does not significantly interact with attention. Indeed, the results of our Bayesian analysis of this effect indicate considerable evidence for the absence of interaction with attention (see Figure 7A). Specifically, 96% of electrodes provided greater evidence for the null hypothesis of no interaction between target type and attention in this time-window (i.e. BF<1), 48% of which provided substantial evidence for the null (i.e. BF<⅓). Within a two-stage model of auditory processing, this effect may be analogous to the mismatch negativity, which has a similar time-course and can be elicited by rare stimuli without attention or conscious awareness (Heilbron & Chait, 2018). However, we had not expected to find evidence for differential processing of matched and mismatched targets during inattention - a result that is seemingly inconsistent with prior evidence that successful comprehension of degraded speech requires top-down expectations (e.g. Sohoglu et al., 2012). Indeed, inattentive participants should have been unable to form top-down expectations that would subsequently elicit a prediction error. One parsimonious interpretation is that the distraction task did not sufficiently direct attention away from the speech stimuli, thus allowing those participants the opportunity to also generate expectations while completing the visual distraction task. However, a Bayesian analysis indicated that our data provide substantial evidence that distracted participants’ memory for the mismatched prime words did not differ from zero. As those words were heard as clear speech, we would expect that memory would be above chance here if the participants were not sufficiently distracted. Therefore, we conclude that the early differential processing of targets during inattention is not the result of insufficient inattention.

**Figure 7.**
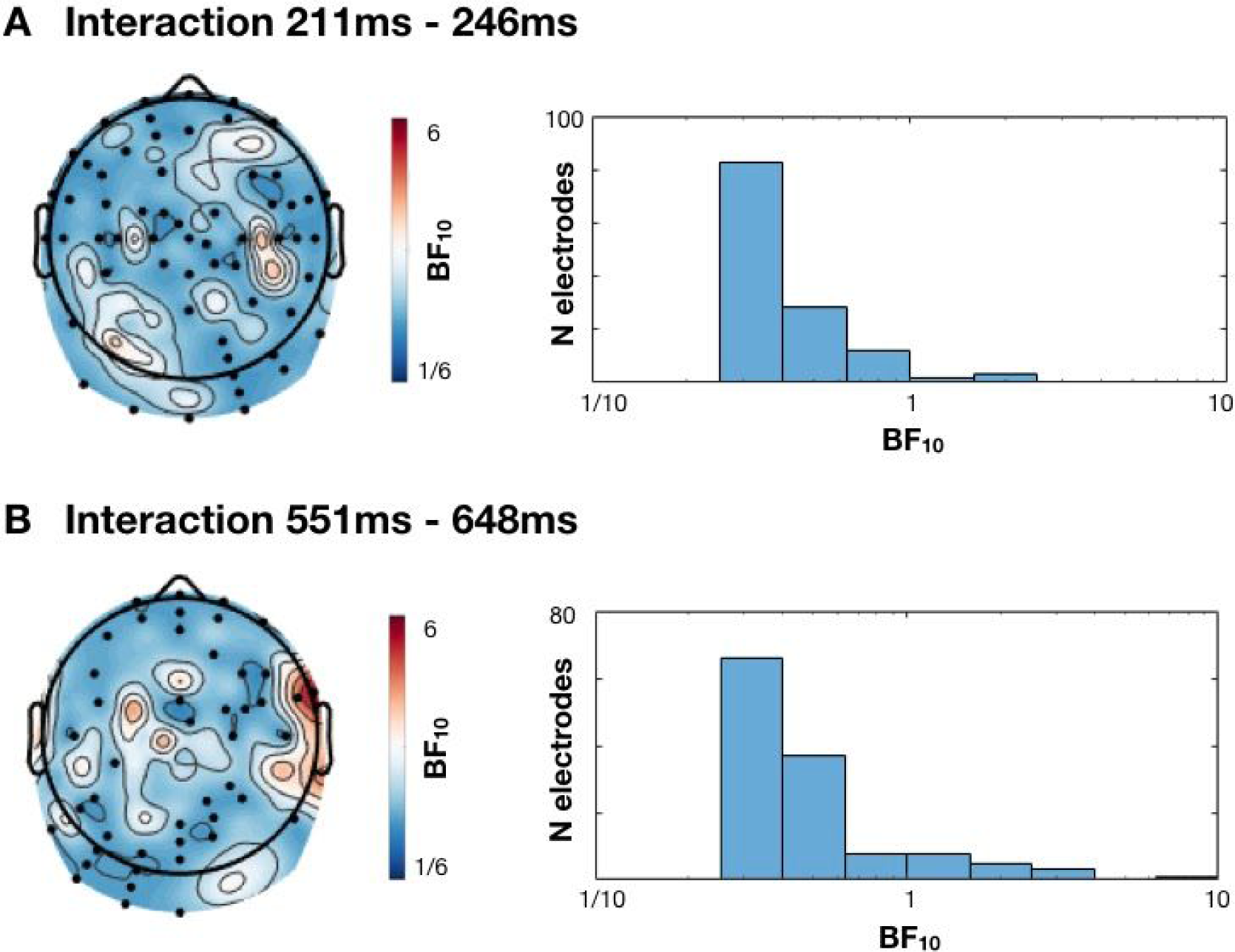
Scalp distribution of Bayes Factors (from Bayesian equivalent t-tests) in tests of the interaction between target type and attention in time-windows [A] 211ms-246ms and [B] 551ms-648ms. All electrodes with Bayes Factors > 3 or < ⅓ are marked on the scalp. Histogram shows the distribution of Bayes Factors across the head.

An alternative interpretation is that the signal reflects the error of a non-conscious expectation that can be generated without top-down influence. For example, previous studies of word-pair priming of noise-vocoded speech have used the written form of the word as the prime stimulus, whereas our prime stimuli were clear versions of the same speech stimulus. Therefore, it is possible that a low-level prediction that an auditory stimulus will have the same envelope as the just-heard auditory stimulus, for example, could be generated inattentively. Indeed, prediction error minimisation accounts posit that expectations are generated at multiple levels of the processing hierarchy. Consequently, while a conscious top-down expectation may not be generated by an inattentive participant, an expectation from a lower level of the hierarchy may nevertheless be instantiated and compared with the sensory input - for example, an expectation that the auditory environment will remain stable, as is one interpretation of the mismatch negativity during inattention (Sussman & Winkler, 2001). Indeed, our source analyses estimate generators of this effect primarily within right temporal lobe and left supramarginal gyrus (see Figure 5C), while a similar previous study involving visual primes (rather than auditory primes as in this study) reported an effect with a similar time-course to be generated within the more canonically semantic regions of left middle and inferior frontal gyri (Sohoglu et al., 2012). We therefore suggest that this effect is a non-semantic error signal, while similar studies that involve visual primes and auditory targets may be more likely to promote semantic expectations. Indeed, the estimated right temporal lobe generator of our error effect is also linked to domain-general processing of complex auditory stimuli, rather than to specific linguistic and semantic processing (McGettigan & Scott, 2012).

As hypothesised, we also observe a later component that is largest for degraded words that ‘pop-out’ into awareness. Specifically, from approximately 250ms to 350ms post-stimulus the ERPs are larger for matched targets than mismatched targets in the attentive group only. Indeed, in that same time window, the ERPs in the distracted group exhibited a similar distribution to the preceding error signal (Figure 3D). The apredictive nature of this ERP component is seemingly at odds with prediction error accounts of evoked potentials. However, the concept of precision is often used to explain such patterns (Kok et al., 2012). Specifically, the error signal is considered to be weighted by the system’s confidence in that signal - its precision. Attention is one mechanism that is thought to increase precision (Hohwy, 2012). Therefore, one could argue that while a fulfilled prediction about an upcoming word elicits little prediction error, an individual’s attention to the word increases precision which multiplicatively leads to a larger precision-weighted prediction error signal (i.e. an evoked potential) than an unpredicted but unattended stimulus. Indeed, the effect of attention to boost the magnitude of evoked potentials is evident in the two main effects of attention we observe prior to this effect (137-207ms and 211-246ms; see Figures 4 and 5). However, it is clear that any observed data can be explained by appealing to the multiplication of two hidden and independently varying signals (namely, error and precision), thus creating issues in rigorously studying the role of precision weighting in perception (cf, Heilbron & Chait, 2018). Nevertheless, under a precision-weighted prediction error interpretation, all evoked potentials should interact with attention, which is demonstrably not the case for our earlier main effect of target (211-246ms; see Figure 7) unless one appeals to complex post hoc interactions of precision and error.

Under a Global Neuronal Workspace interpretation, this later component could be considered to reflect the breakthrough of a stimulus representation into conscious experience (Alsufyani et al., 2019). While we did observe weak evidence of recollection of mismatched targets (p=.020; BF10 = 2.720), indicating that mismatched targets were not entirely unintelligible perhaps due to a degree of perceptual learning across the experiment (Hervais-Adelman et al., 2008), attentive participants’ ratings of intelligibility (noisiness) were entirely consistent with the pop-out of meaning following matched primes (Davis et al., 2005). Furthermore, this component was larger for subsequently remembered items than for subsequently forgotten items, albeit weakly (p=.049; BF10=1.628). The link to subsequent successful recognition provides further evidence to link this ERP component with a conscious experience on the part of the listener. However, this effect is earlier than we predicted based on a typical two-stage profile and its scalp distribution is more reminiscent of a P3a than the P3b or other late positive components typically linked to global-workspace breakthrough effects. Nevertheless, P3a-like components have been observed in breakthrough contexts (Bowman et al., 2013) and Global Neuronal Workspace theory broadly posits that scalp positivities, as observed here, reflect the ignition of a representation into conscious access (Dehaene & Christen, 2011). Source estimates of other late positivities within this framework typically involve generators distributed across the cortex, consistent with a brain-wide ignition into conscious access (e.g. Bekinschtein et al., 2009). However, while the source estimate of our observed pop-out effect includes some weak evidence of generators distributed across lobes and hemispheres (see Figure 3C), the focus is estimated to be in the right middle temporal gyrus and right fusiform gyrus. Interestingly, the late positivity described by Rohaut et al. (2015), and linked to the stage of conscious access of meaning within the two-stage profile, was also estimated to be generated within right fusiform gyrus, as well as left dorsolateral frontal cortex. Furthermore, there is evidence for greater activity within the right fusiform gyrus when the meaning of speech is task-relevant (von Kriegstein et al., 2003). We conclude therefore that our ERP positivity reflects conscious access of the meaning of speech. While we cannot rule out the potential role of task-related post-perceptual processing rather than conscious access itself (Aru et al., 2012), we argue that the majority of evidence for components linked to such processes occur later in time than the pop-out effect observed here (i.e. after ∼350ms; e.g. (Pitts et al., 2014; Schelonka et al., 2017).

In a later time-window, from approximately 550ms to 650ms, we also observed more extreme ERPs for mismatched targets than for matched targets that did not interact with attention (see Figures 6 and 7). The scalp distribution of this effect is markedly similar to that reported in an overlapping time-window of 450ms to 700ms post-target by Sohoglu et al. (2012); see Figure 4B in that paper) who also found that magnetoencephalography (MEG) sensor data in the same time-window significantly predicted trial-by-trial ratings of speech clarity, such that reduced neural responses to matched targets were accompanied by increased experiences of speech clarity. Our source estimates indicated a primarily right posterior superior temporal generator for this effect, while Sohoglu et al. (2012), with more sensitive MEG source analyses, report right temporal generators alongside bilateral inferior frontal and middle occipital gyri. Sohoglu et al. (2012), therefore, conclude that this effect reflects the neural processes that generate the experience of speech clarity. On that basis, we would expect this effect to interact with attention in this study. However, we find no evidence for this interaction. Nevertheless, while Bayes equivalent t-tests at each electrode in this time-window indicated substantial evidence for no interaction in the majority of electrodes, two electrodes did exhibit substantial evidence for an interaction (i.e. BF10>3; Figure 7B). As our cluster forming threshold required 4 neighbouring electrodes, it is possible that this effect does indeed interact with attention, but to an extent that is not evident with our specific analysis choices. If that were the case then, these later effects may indeed reflect processes associated with the conscious experience of meaning, or may reflect consequent processes such as those in service of task demands - i.e. providing a judgment of the noisiness of the stimulus (Aru et al., 2012).

## Conclusions

Our results indicate a link between the conscious experience of semantic pop-out in comprehension of degraded speech and a positive-going ERP in the range of 300ms post-stimulus - consistent with a Global Neuronal Workspace framework. Prior to this positivity, ERPs appear to reflect the error of non-semantic predictions, consistent with prediction error minimisation accounts. To consider our observed late positivity within the same prediction error account requires a post hoc appeal to freely varying precision weighting that is not straightforwardly verified. We therefore suggest that our data are consistent with early negative-going ERPs as reflections of prediction error while later positive-going ERPs reflect conscious access and processes in support of task demands (e.g. Dehaene & Christen, 2011; Rohaut et al., 2015).

## Acknowledgments

This work was supported by generous funding from the Medical Research Council IMPACT Doctoral Training Programme at the University of Birmingham (Scholarship to LB) and a Medical Research Council New Investigator Research Grant (MR/P013228/1; Principal investigator: DC).

## Data availability

All data and scripts are available at https://osf.io/pckyv/

## Supplementary Materials

**Supplementary Table 1:**
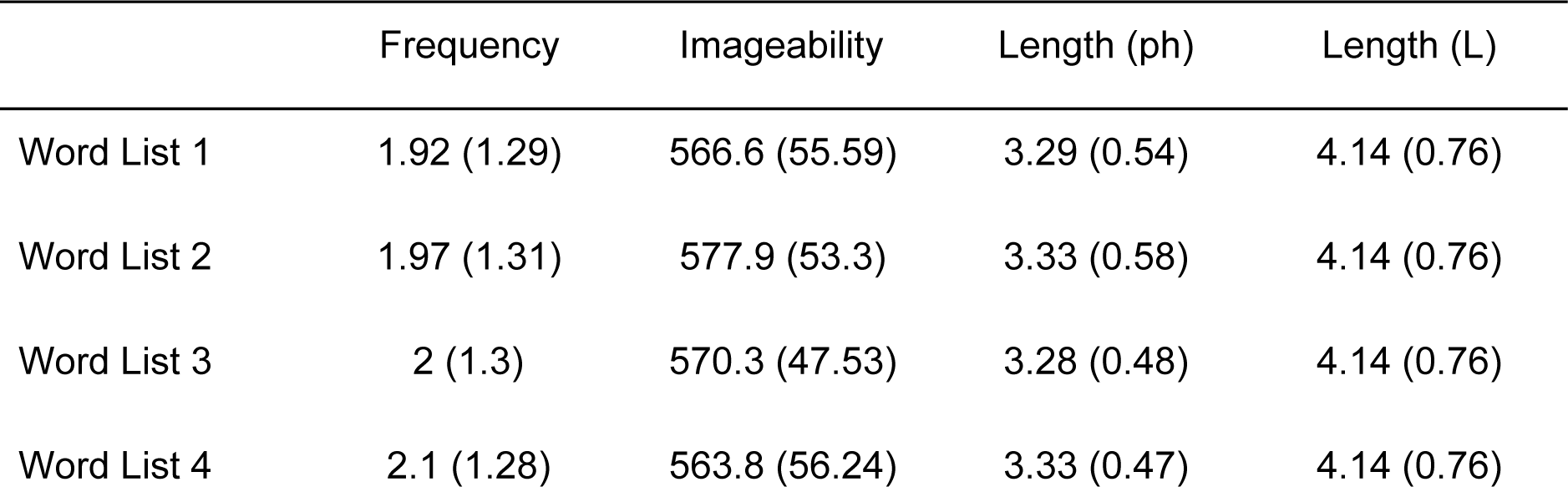
Mean and standard deviation (in brackets) of the word list characteristics: frequency (ln[BNC]), imageability, length in phonemes, and length in letters.

|  | Frequency | Imageability | Length (ph) | Length (L) |
| --- | --- | --- | --- | --- |
| Word List 1 | 1.92 (1.29) | 566.6 (55.59) | 3.29 (0.54) | 4.14 (0.76) |
| Word List 2 | 1.97 (1.31) | 577.9 (53.3) | 3.33 (0.58) | 4.14 (0.76) |
| Word List 3 | 2 (1.3) | 570.3 (47.53) | 3.28 (0.48) | 4.14 (0.76) |
| Word List 4 | 2.1 (1.28) | 563.8 (56.24) | 3.33 (0.47) | 4.14 (0.76) |

**Supplementary Table 2.** ANOVAs and equivalent Bayesian ANOVAS testing for differences across lists at each word characteristic.

|  | F | *p* | BF10 |
| --- | --- | --- | --- |
| Frequency | 0.233 | 0.873 | 0.021 |
| Imageability | 0.779 | 0.507 | 0.054 |
| Length (ph) | 0.217 | 0.885 | 0.021 |
| Length (L) | <.001 | 1 | 0.016 |

**Supplementary Figure 1:**
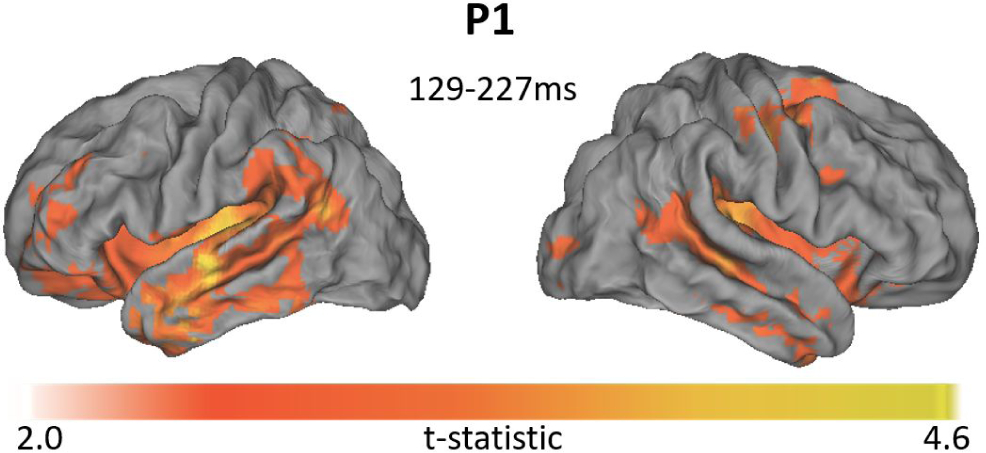
P1 source estimation validity check. As a validity check of our source estimation method, we estimated the sources of the auditory P1 component. Specifically, we calculated the global field power of the grand average of all EEG trials to identify the boundaries of the first component. The figure shows a bilateral temporal focus of the estimated sources of this component (129-227ms), broadly consistent with temporal lobe generators of early auditory ERPs.

